# Atomic-scale Quantum Chemical Calculation of Omicron Mutations Near Cleavage Sites of the Spike Protein

**DOI:** 10.1101/2022.06.03.494698

**Authors:** Puja Adhikari, Bahaa Jawad, Rudolf Podgornik, Wai-Yim Ching

## Abstract

The attachment of the Spike-protein in SARS-CoV-2 to host cells and the initiation of viral invasion are two critical processes in the viral infection and transmission processes in which the presence of unique furin (S1/S2) and TMPRSS2 (S2’) cleavage sites play a pivotal role. In this study, we provide detailed analysis of the impact of the BA.1 Omicron variant mutations, vicinal to these two cleavage sites using a novel computational method based on Amino acid – amino acid bond pair unit (AABPU), a specific protein structural unit in 3D as a proxy for quantifying the atomic interaction. We have identified several key features related to the electronic structure as well as bonding of the Omicron mutations near the cleavage sites that significantly increase the size of the relevant AABPUs and the fraction of the positive partial charge. These results of the ultra-large-scale quantum calculations enable us to conjecture on the biological role of Omicron mutations and their specific effects on cleavage sites, as well as identify the principles that can be of some value in analyzing other new variants or subvariants.

## 1. Introduction

SARS-CoV-2 virus continues to mutate and evolve, leading to major variants of concern (VOC) that can significantly change the virus characteristics [1]. These VOC result in increased infectivity, transmissibility, and severity as well as reduced efficacy of available therapies [1–3]. The Omicron variant (OV), the most recently identified VOC, rapidly became the dominant strain globally due to an unprecedented number of mutations in the spike (S) protein, making it highly contagious and/or vaccine-resistant [4–6]. During the early stages of virus infection, the S-protein mediates both the receptor binding *via* its S1 subunit as well as the membrane fusion *via* its S2 subunit. Upon binding of the S-protein to the human angiotensin-converting enzyme 2 (ACE2) in the host cell, the S-protein is proteolytically cleaved at two protease recognition sites: the S1/S2, or furin cleavage site, and the TMPRSS2 S’, or *transmembrane serine protease 2 cleavage site*, in order to activate the fusion machinery [7]. The furin S1/S2 site is located at the boundary between the S1 and S2 subunits, exhibiting a unique polybasic insertion furin recognition site _681_PRRAR|S_686_ (“|” denotes proteolytic cleavage site) [8], while the TMPRSS2 S’ site is located just upstream of the fusion peptide (FP) domain of the S2 subunit. These cleavage sites play an essential role in viral infectivity, transmissibility, fusogenicity and pathogenicity [8,9]. Mutations at the cleavage sites have been shown to promote more efficient cell–cell fusion in both Alpha and Delta VOCs and can facilitate and enhance cell entry thus increasing transmissibility [10,11]. Near the S1/S2 cleavage site, OV has the same mutation of P681H as the Alpha variant, connected to the increased infectivity, in addition to two other mutations of H655Y and N679K [12]. Moreover, OV has six unique mutations at the S2 subunit (N764K, D796Y, N856K, Q954H, N969K, L981F) that have not been previously detected and whose biological functions are as yet unknown [12]. Therefore, it is urgent to identify the role of these novel OV mutations and investigate their impact on the S-protein, particularly around the proximal region of the cleavage sites. This will provide information on the change in interatomic interaction and guide effective therapeutic strategies to overcome current and future cross-strain SARS-CoV-2 infections.

S-protein of SARS-CoV-2 is the main target of most therapies [13–16]. It appears in the trimeric form, with each protomer comprising two functional subunits, S1 and S2, that are demarcated by a furin cleavage site (S1/S2), as shown in **Figure 1**. S1 contains the signal peptide (SP), N-terminal domain (NTD), receptor-binding domain (RBD), subdomain 1 (SD1), and subdomain 2 (SD2), while S2 consists of fusion peptide (FP), heptad repeat 1 (HR1), central helix (CH), connector domain (CD), heptad repeat 2 (HR2), transmembrane domain (TM), and cytoplasmic tail (CT) [17]. Viral entry into the host cell depends on these two subunits. The cleavage sites play a role in facilitating SARS-CoV-2 entry. The cleavage activation mechanism occurs at S1/S2 and S2’ is a very complex process [18]. Briefly, after RBD of S1 subunit recognizes and attaches the ACE2 receptor, the S-protein is initially cleaved at the S1/S2 boundary by the furin protease, in which both S1 and S2 remain non-covalently associated in the prefusion conformation [7,18]. This is followed by a second cleavage at the S’ site with the S protein undergoing significant conformational changes, resulting in the dissociation of S1 and the irreversible refolding of S2 into a post-fusion structure [19]. This causes the virus and host cell membranes to fuse, allowing infection to begin. Due to the complexity of the role played by the cleavage sites in virus transmission and the lack of information on this process of Omicron variant, accurate state-of-the-art *ab initio* computational study could reveal some of intricate details related to the amino acid mutations close to the cleavage sites at the molecular, amino acid and atomic levels. Such critical information is currently missing.

**Figure 1.**
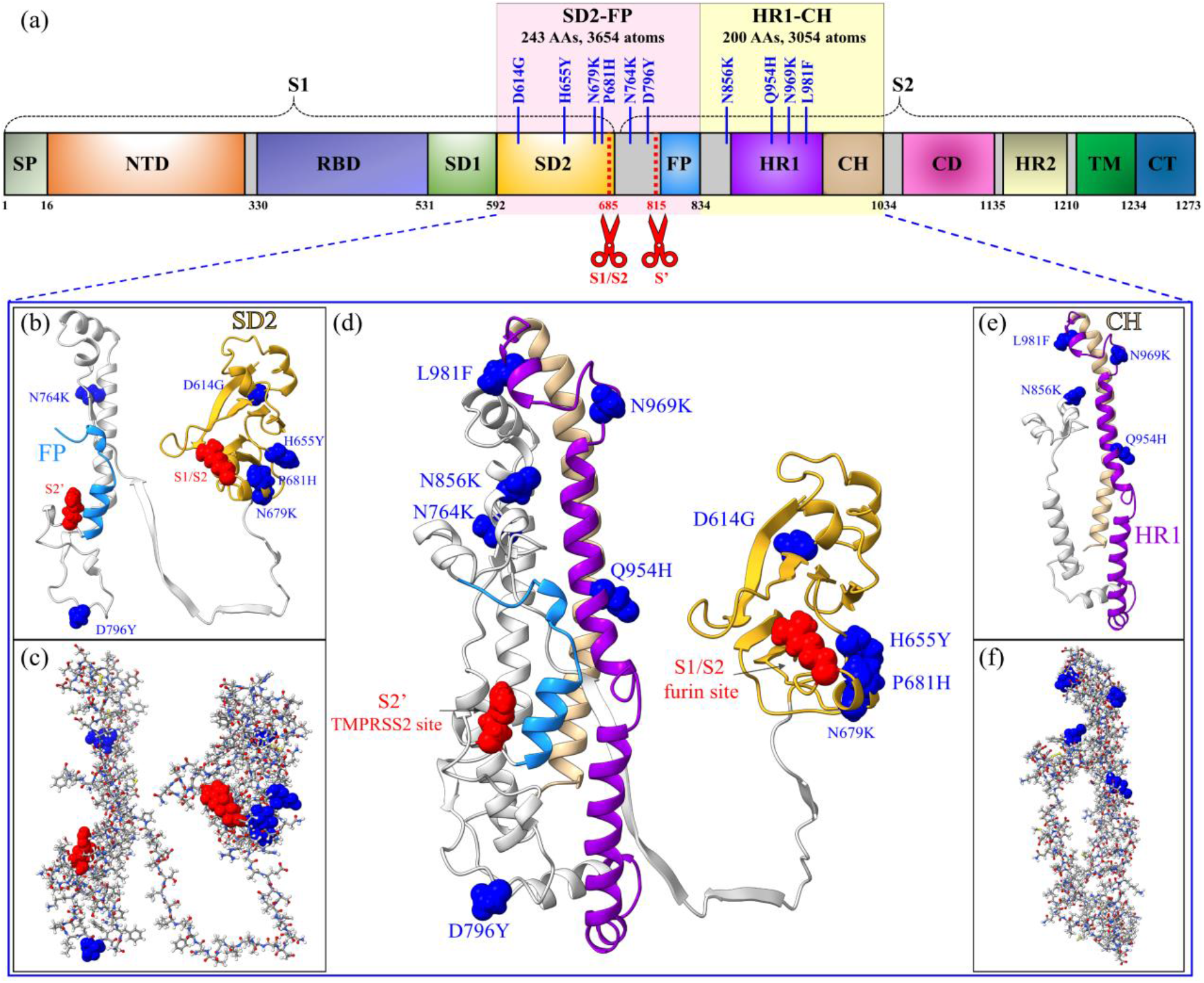
Graphic illustration of SD2-FP and HR1-CH models. (a) Schematic of S-protein primary structure divided into domains with cleavage sites S1/S2 and S2’ denoted by scissors and highlighting the SD2-FP and HR1-CH regions with their ten Omicron mutations. (b) and (c) SD2-FP model in ribbon and ball and stick representations respectively. Their six Omicron mutations are marked and shown by a blue sphere, while the S1/S2 and S2’ cleavage sites are represented by a red sphere. (d) Ribbon figure of SD2-FP and HR1-CH showing the two cleavage sites (S1/S2 and S2’) and marked the 10 Omicron mutations in these regions. (e) and (f) HR1-CH model with their 4 Omicron mutations in different representations. The red, blue, grey, yellow, and white in ball and stick figures are for O, N, C, S, H atoms respectively.

In this work, we focus on the impact of the Omicron mutations close to the cleavage sites S1/S2 and S2’ between SD2 to FP (SD2-FP) and HR1 to CH (HR1-CH), which contains 10 mutations (D614G, H655Y, N679K, P681H, N764K, D796Y, N856K, Q954H, N969K, L981F) (see **Figure 1**). Most importantly, the last six mutations (N764K to L981F) in S2 are the unique mutations in OV. We constructed two models for SD2-FP and HR1-CH, one for the wild-type (WT) or unmutated case and the other one for the mutated OV. **Figure 1(d)** shows the ribbon structure of OV from the SD2-FP and HR1-CH models, which contain the 2 cleavage sites S1/S2 and S2’ as well as the 10 mutated AAs, as marked. The two systems must be calculated separately within the ultra-large first principles *ab initio* approach for biomolecules, based on the *divide and conquer strategy* [20,21]. The SD2-FP model has 3654 atoms (WT) and 3681 atoms (OV), and HR1-CH has 3054 (WT) and 3071atoms (OV), including H atoms. **Figure 1(d)** illustrates the ways to join the two parts seamlessly as will be elaborated in **Section 4.1**. In this way, we can assess the effects of the 2 cleavage sites in relations to the mutation effect in OV much more efficiently.

The purpose of this paper is to better understand the role of the S-protein polybasic cleavage sites, S1/S2 and S2’ of WT or OV, and to show how the Omicron mutations in the vicinity of these sites impact their interatomic interactions in the S-protein. Specifically, this study provides detailed computational results to assess the role of each of the 10 OV mutations in SD2-FP and HR1-CH models to ascertain the possible reasons for the rapid infection rate of OV. We employ the novel concept of amino acid-amino acid bond pair (AABP) as specific protein structural unit [22] to quantify and characterize the details of the impact of OV mutations which could help to determine their role in high infectivity.

## 2. Results

### 2.1 Amino acid amino acid bond pair (AABP) unit

In the Materials and Methods Section, we will describe the previously introduced AABP [23] as an extension of the AA-AA interaction description, that includes also the contribution from non-local AAs that are not vicinal along the primary AA sequence. The AABP considers all possible bonding between two AAs including both the covalent and the hydrogen bonding (HB). This single quantifier, derived directly from *ab initio* quantum chemical calculation of the electronic structure in large supercells of several thousands of atoms, reflects the internal bonding strength between all relevant amino acids. Such information is essential for analyzing the effects of mutations on the properties of a protein as well as the whole virus. AABP can be further resolved into nearest neighbor (NN) bonding and non-local (NL) bonding from non-NN pairs along the protein sequence. We can also single out the contribution to the total AABP from HB. In this sense AABP is an ideal parameter to characterize interactions between different AAs or groups of interacting AAs in biomolecules, *i*.*e*., we can consider the AABP to characterize a specific biological unit: the *AABP unit (AABPU)*. In reference [22], AABPU has been thoroughly described and successfully used to investigate the ten mutations at the interface between the receptor domain motif (RBM) and the human ACE2 receptor of the Omicron variant. Hence, in terms of the interactions the AABPU can be considered as distinct structural units in proteins, capable of providing full physical insight into properties such as interatomic interactions, partial charge distributions, size, shape, NL-AA interactions etc. This approach furthermore allows us to consider more relevant details of biological structure than simply the AA sequence in the case of proteins. The results for the structure and some pertinent properties of the ten mutations in the two models containing the cleavage sites are summarized in **Table 1**. WT (wild type) stands for the unmutated type and OV stands for the mutated Omicron variant (OV). To facilitate discussion, these 10 mutations are divided into 3 groups separated by the cleavage site S1/S2 and S’ indicated by the red dashed lines in the **Table 1**.

**Table 1.** AABP results due to mutation Omicron Variant close to cleavage site, six SD2-FP region and four HR1-CH regions. Colored dash lines are locations of S1/S2 and S2’cleavages separating the 10 mutations. We label the first 6 mutations as group A, the second 2 mutations as group B and the last 4 mutations as group C to facilitate the association of their relative positions with respect to the two cleavage sites. AABP is in unit of electrons (e-).

| Models | Total AABP | NN AABP | NL-AABP | AABP (HB) | # NL AAs | Vol (Å<sup>3</sup>) | Surface (Å<sup>2</sup>) | PC* (e-) |
| --- | --- | --- | --- | --- | --- | --- | --- | --- |
| WT D614 | 0.917 | 0.912 | 0.005 | 0.040 | 5 | 745.2 | 644.6 | 0.055 |
| OV G614 | 0.908 | 0.907 | 0.002 | 0.042 | 3 | 640.3 | 564.5 | 0.961 |
| WT H655 | 0.976 | 0.968 | 0.007 | 0.032 | 5 | 909.3 | 727.8 | -0.827 |
| OV Y655 | 0.971 | 0.965 | 0.006 | 0.032 | 5 | 944.5 | 745.3 | -0.782 |
| WT N679 | 1.022 | 0.956 | 0.066 | 0.081 | 3 | 624.6 | 533.9 | 0.833 |
| OV K679 | 1.011 | 0.963 | 0.048 | 0.072 | 4 | 807.4 | 657.4 | 1.766 |
| WT P681 | 1.117 | 1.064 | 0.054 | 0.064 | 5 | 872.2 | 757.9 | 2.862 |
| OV H681 | 1.032 | 0.984 | 0.048 | 0.068 | 5 | 933.9 | 832.1 | 3.881 |
| WT N764 | 1.130 | 1.008 | 0.121 | 0.137 | 6 | 959.5 | 775.1 | 1.043 |
| OV K764 | 1.118 | 1.019 | 0.100 | 0.118 | 8 | 1211 | 986.4 | 1.181 |
| WT D796 | 1.175 | 1.124 | 0.052 | 0.066 | 2 | 527.3 | 476.6 | 0.043 |
| OV Y796 | 1.051 | 1.000 | 0.051 | 0.070 | 2 | 599.4 | 519.3 | 1.040 |
| WT N856 | 0.935 | 0.894 | 0.041 | 0.069 | 3 | 609.9 | 554.0 | 0.783 |
| OV K856 | 0.937 | 0.902 | 0.036 | 0.065 | 6 | 1081.0 | 822.5 | 1.801 |
| WT Q954 | 1.148 | 1.008 | 0.140 | 0.152 | 7 | 1063.0 | 822.3 | 0.028 |
| OV H954 | 1.146 | 1.008 | 0.139 | 0.155 | 7 | 1077.0 | 813.9 | 0.000 |
| WT N969 | 0.938 | 0.907 | 0.031 | 0.052 | 5 | 774.1 | 624.5 | 0.188 |
| OV K969 | 0.946 | 0.913 | 0.033 | 0.053 | 6 | 893.1 | 675.4 | 0.829 |
| WT L981 | 0.898 | 0.893 | 0.005 | 0.036 | 5 | 960.1 | 754.9 | 0.090 |
| OV F981 | 0.917 | 0.888 | 0.029 | 0.059 | 4 | 811.7 | 677.4 | 0.184 |

The salient features of the results in **Table 1** are summarized as follows:

1. The largest total AABP is from the site 796 close to cleavage site S2’ with only 2 NL-AAs. D796Y mutation reduces AABP slightly, but it increases size, surface exposure, and positive charge. The largest NN-AABP is from the same site 796 with only 2 NL-AAs. This confirms that *NN AAs is the dominant interaction* as evidenced by the primary sequence of the S-protein.
2. The smallest total AABP comes from site 981, which is relatively far from S’ cleavage site but near the prefusion-stabilizing two-proline (2P) mutations (K986P and V987P) utilized in Pfizer-BioNTech and Moderna vaccines [13,14]. Site 981 also has the smallest NN-AABP in WT but with NL-AAs of 5.
3. The 954 site has the largest NL-AABP contribution among all ten mutations. It is part of the HR1 domain, as are the other two sites, 969 and 981, which have low NL-AABPs. These three Omicron mutations have slightly different interatomic interactions than their WT counterparts, while their size, shape, and PC*s are all changed. We speculate that these mutations may play a role in the binding of HR1 and HR2 domains to enhance the formation of six-helix-bundle (6-HB), which brings the viral lipid and the host lipid membranes close together, resulting in membrane fusion and initiation of infection [24]. Here, it should be mentioned that HR1-CH model alone is insufficient to assess this binding process. All atoms of the post-fusion S-protein must be included, which is currently impossible to do in a single *ab initio* calculation.
4. The smallest NL-AABP is from site 614 in group A ahead of the S1/S2 cleavage site. D614G mutation reduces the number of NL-AAs from 5 to 3 but results in a shift in charge distribution toward a more positively charged state, which could enhance the susceptibility of protease cleavage at the S1/S2 junction and/or promote the up conformation of the S-protein, as previously reported [25–27].
5. The largest contribution from HB to total AABP is from the site 954 with the largest NL-AABP, while the smallest HB contribution is from sites 614 and 655 in group A and with concomitant small NL-AABP. This attests to the importance of the contribution of HB to the overall bonding network.
6. The role of the PC* distributions is obvious from five mutations D614G, N679K, P681H, D796Y, and N969K. They exhibit a significantly changed, more positive, PC*. Importantly, the mutation at 681 site, which is adjacent to the furin cleavage site, has been reported to play a significant function in the cleavage process [10,11,21,28]. Increasing the positive charge of P681H is necessary for the host furin-like proteases to cleave the S-protein [11]. Additionally, N679K mutation is also located in the furin cleavage region and has been reported to increase the furin-mediated cleavage of Omicron [29]. However, some investigations have indicated that these N679K and P681H mutations do not enhance the S-protein cleavage processing and may even be less efficient [12,30–32]. This suggests that additional mutations near the furin cleavage site may interfere with its cleavage.

These observations clearly indicate that the changes in the total AABP values and their non-local components depend on the nature of the substitution, interatomic interactions, location with respect to the cleavage sites, and the HB contribution. The HB network between biological units controls to a large extent how they interact, which may affect how the S-protein interacts with the host cells or escapes from antibodies, based on the possible changes in the structural geometry to be discussed later.

In **Figure 2**, we plot the main results of **Table 1** with the histogram bars, showing the changes due to mutations side by side. We have also added vertical red dashed lines like the horizontal dashed red lines in **Table 1. Figure 2** essentially reflects the observations listed above. The distribution of the total AABP values in (a) is closely mimicked by the NN-AABP values in (b) reflecting fact the AA sequence is always the most relevant ordering in proteins but the addition of NL-AABP is not negligible as shown in (c) and accounts for the majority of HB contributions to the total AABP as illustrated in (d). An important accomplishment is that we can actually calculate the volume and surface areas of each AABPU as a specific structural unit. Another important observation from **Figure 2** is that mutation slightly decrease the AABP values in most cases, but they tend to increase the volume of AABPU with some of them rather substantially (N764K, N856K). The plots for the AABPU volume and surface area in (e) and (f) mimic each other as expected but do not coincide, since the complexity of the shapes of the AABPU plays a role that defies a simple quantification. This will be illustrated in **Figure 3** and **Figure 4** to follow. Out of 10 mutations only 2 have volume decreased or 20%, D614G at far left and L981F at far right.

**Figure 2.**
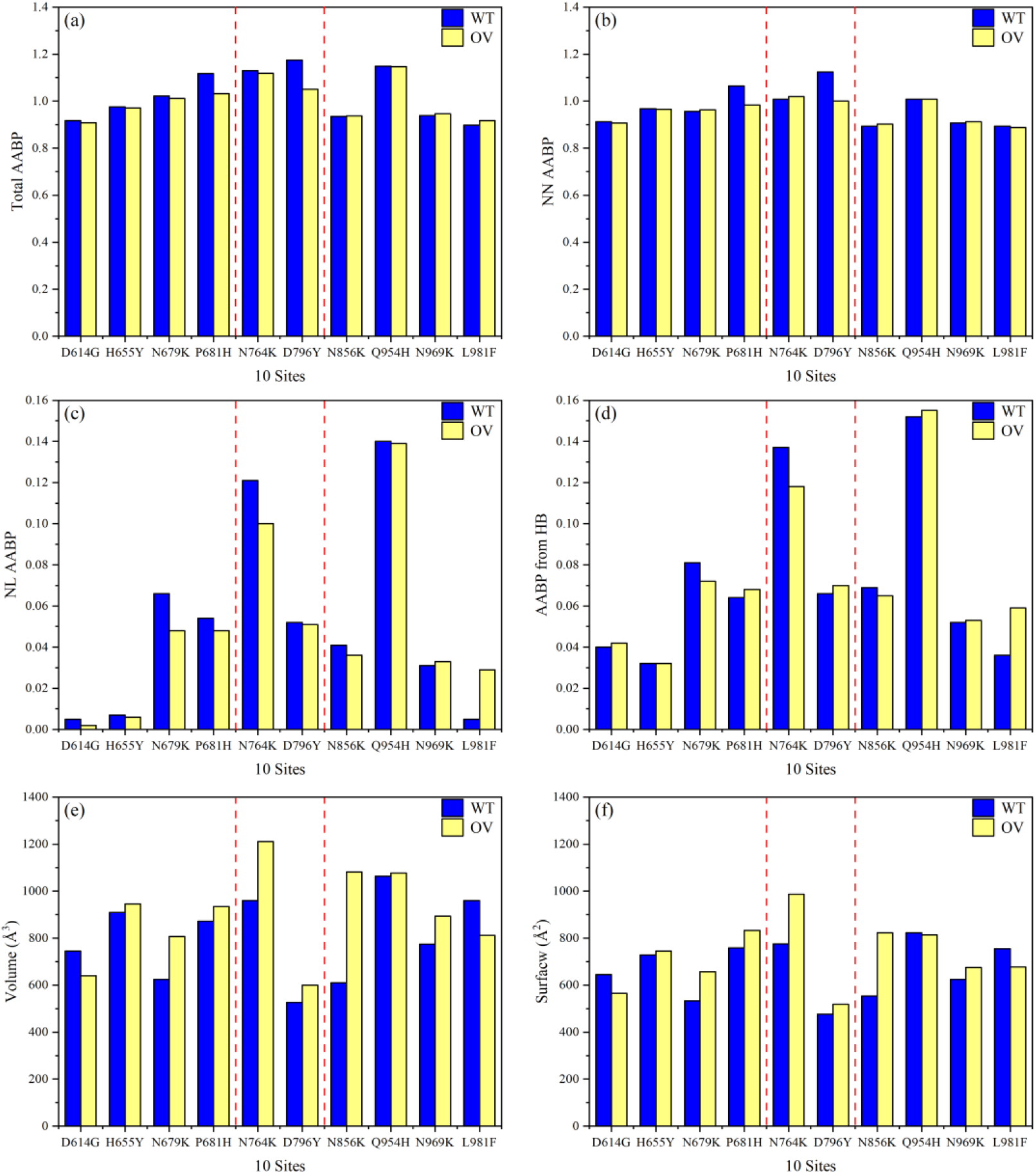
Comparison of (a) Total AABP, (b) NN AABP, (c) NL AABP, and (d) AABP from HB, for 10 unmutated (WT) and mutated (OV) AAs. (e) Volume, and (f) Surface.The two dashed red lines shows the two cleavage sites.

In addition to revealing the details on the 10 Omicron mutations, we can vividly depict the size and shape for each of the AABPU (see **Figure 3**). For the benefit of a compact illustration, the geometrical scale used in **Figure 3** is kept the same. The 10 mutations are separated into three groups A, B and C as in **Table 1. Figure 3** displays mutation effect in 2D plots for the 3D structures with the same length scale. The volume and surface data in **Table 1** provide the geometry of the Omicron pictorially by connecting the mutation effect with the proximity to the cleavage locations. With exception of D16G and L981F, OV mutations tend to increase their volume. N856K, which is located between FP and HR1 domains within group C, has the greatest change in volume. The volume (surface) increases by 77.2% (48.5%), with significant changes in the shape of the AABPU. While the smallest change in volume comes from Q956H. There are two mutations, D614G and L981F reduced their volume. The smallest change in other 8 mutations is the increased volume in Q956H.

**Figure 3.**
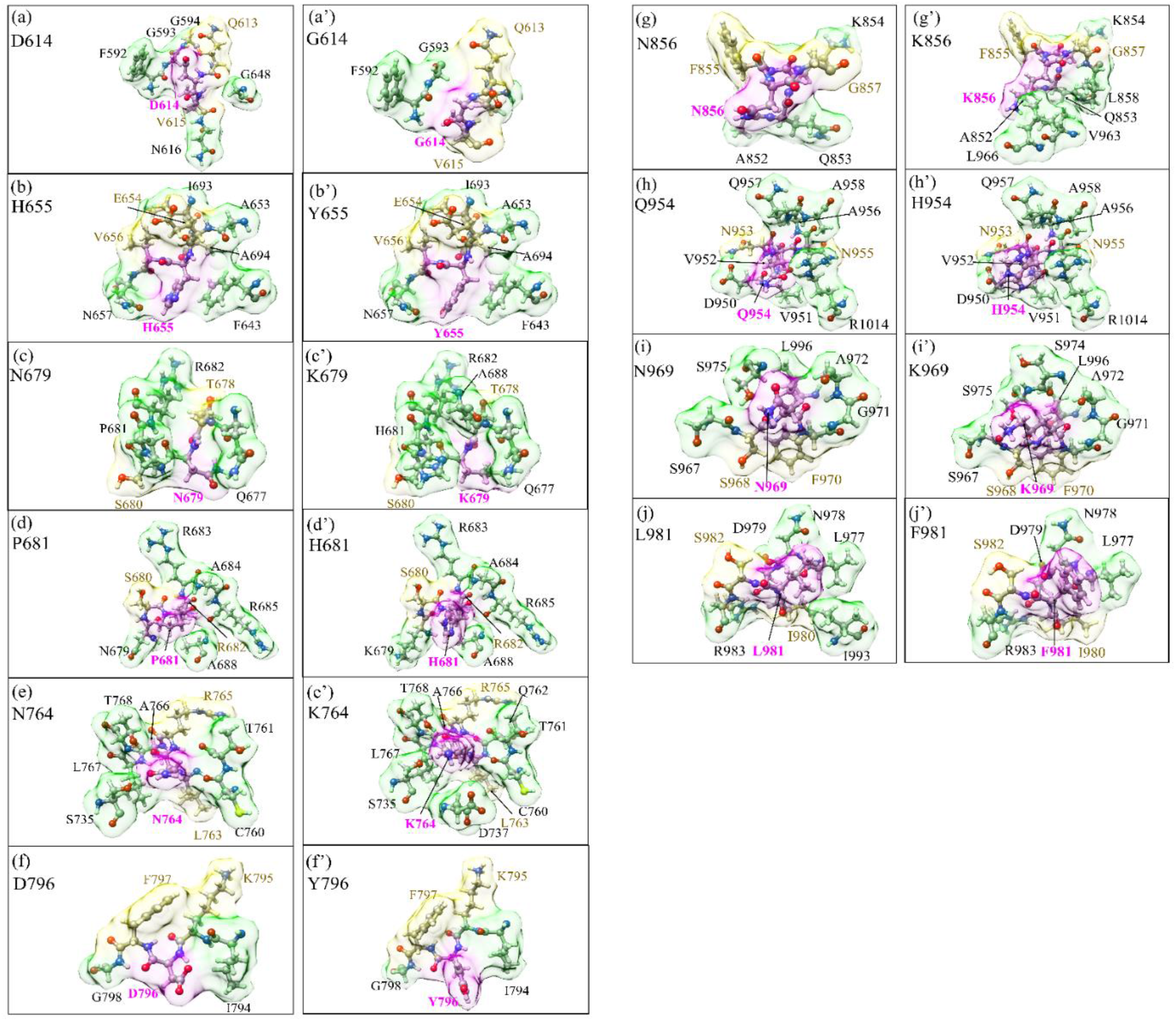
Distribution of the six mutation sites in SD2-FP (left panel) and 4 mutation sites in HR1-CH (right panel). Within each panel, (a) to (j) on the left column is for the WT, and (a’) to (j’) on the right column is for the OV. The surface of mutated sites is shown in magenta, surface of NN and NL are shown in yellow and green respectively. All NN and NL AAs are marked near to their surface in brown and black respectively.

To better illustrate the size and shape due to mutation, we replot the cases of N856K and Q954H mutations in more details in **Figures 4(a)** and **(b)** respectively with corrected length scale and in three different orientations. As shown in the **Table 1**, N856K has the largest increase in volume of 77.2% and the shape of the AABPU is drastically different between WT and OV. On the other hand, Q954H has the largest volume both before and after mutation, so the difference in the increased volume is small. As expected, in Q954H there is very small change in the shape of its AABPU before and after mutation with a tiny increase of 1.3%. To further demonstrate the power of our graphical illustration, we plot similar figures for N764K and D796Y mutations in **Figures 4(c)** and **(d)**, respectively. Both N764K and D796Y are between the two cleavage sites S1/S2 and S2’ at the N-terminus of the S2 subunit and both increased their volume by 26.2% and 13.7% respectively. Moreover, N764 has the largest number of NL-AAs, increasing from 6 to 8 upon mutation. D796 has the fewest NL-AAs of 2 and there is no change in this number after mutation. The biological function of the D796Y mutation is still unclear. Previously, it has been revealed that the mutation at this site impairs the neutralization susceptibility of antibodies and reduces SARS-CoV-2 infectivity [33]. Based on detailed *ab initio* calculations, we expect that shifting the partial charge of the N764K and D796Y toward more positive values as their sizes increase could impact the cleavage site S’, the interactions between S1 and S2 subunits, or the susceptibility to neutralizing antibodies. Further investigations are necessary here. Additional figures for these two special cases provide us some new insights on the complex interplay of the proximity to the cleavage sites.

**Figure 4.**
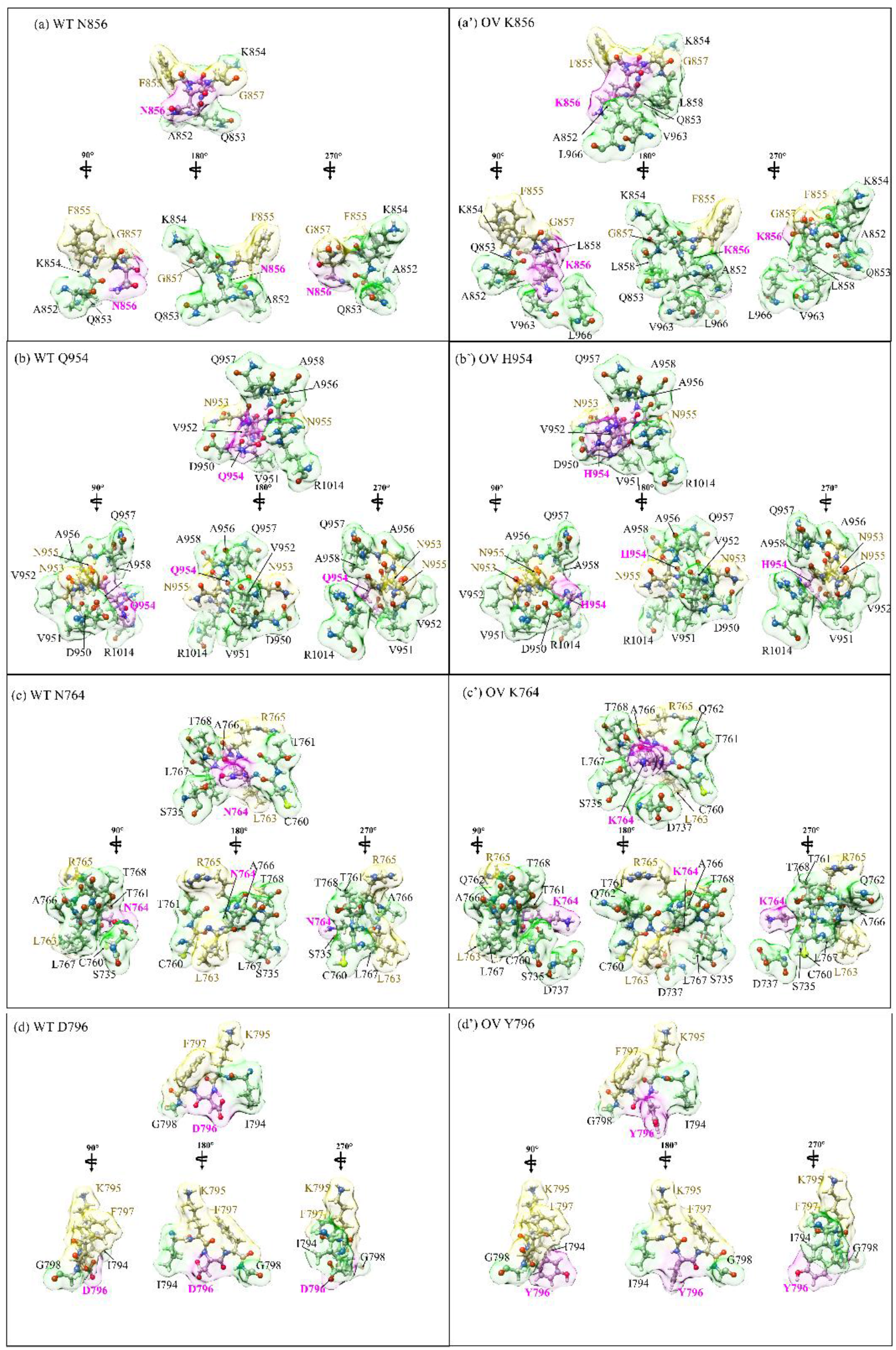
Changes in intramolecular shape and size of specific mutations in the Omicron variant. Surface figures in different orientation for N856K in (a) and (a’), Q954H in (b) and (b’), N764K in (c) and (c’) and D796Y in (d) and (d’). The WT on the left while OV on the right. The surface of mutated sites is shown in magenta, surface of NN and NL are shown in yellow and green respectively. All NN and NL AAs are marked near to their surface in brown and black respectively.

### 2.2 Electronic structure

The simplest way to demonstrate the electronic structure is by presenting the total and partial density of states (TDOS and PDOS), a representation commonly adopted in condensed mater physics and/or materials science. In small molecules, they are usually presented in the form of energy levels close to the highest occupied molecular orbital (HOMO) and the lowest unoccupied molecular orbital (LUMO) separated by a small gap. This is obviously unpractical for large biomolecular systems with up to hundred thousand or more energy levels. In **Figure S1(a)** and **(b)**, we plot the TBOD of the two models SD2-FP (3654 atoms (WT) and 3681 atoms (OV)) and HR1-CH (3054 atoms (WT) and 3071 atoms (OV)) in each frame. The result is very interesting. For SD2-FP, the peaks in the TDOS are very close to each other with those OV at lower energy by about 0.15 eV and with a clear HOMO-LUMO gap of 1.60 eV. For SD2-FP, the structure of WT and OV are still very similar but those for OV are shifted to a lower energy with the same HOMO-LUMO gap of 1.60 eV. We set the energy scale of 0. eV or the HOMO level of WT for easy comparison. For HR1-CH, the general features are similar to SD2-FP except the peak separations are larger, of 0.71 eV. The HOMO-LUMO gap is 1.40 eV for WT and 1.70 eV for OV. What **Figure S1** shows is that the overall features of TDOS in WT and OV are quite similar. But the TDOS in OV is lower in energy by about 1.5 eV from WT in HR1-CH compared to SD2-FP. Whether this feature is related to the relative positions of the cleavage sites S1/S2 and S2’ is not clear at this time.

The TDOS in **Figure S1** is resolved into PDOS for each mutation in **Figure S2(a)** and **(b)**. While the peak positions in PDOS for WT and OV are still very similar, there are some interesting mutation-dependent features. For example, the area of PDOS below HOMO which accounts for the number of electrons is larger in OV than in WT, except for mutation D614G in SD2-FP model. This is consistent with the fact that these mutations increase the volume of AABPU with larger number of AAs, except in the case of mutation D614G. Although the mutation L981F in HR1-CH also decreases its volume by a small amount (see **Table 1**), so are the small difference in the area below HOMO in **Figure S2(b)**. Both these two mutations are farther away from the cleavage sites.

### 2.3 Interatomic bonding

Figure 5. shows the distribution of bond order (BO) vs bond length (BL) for all atomic pairs in the models SD2-FP and HR1-CH with BL ranging from (a) 0.5 Å to 2.0 Å and (b) 2.0 Å to 4.5 Å. The WT and OV are shown in open and closed symbols respectively, for all the atoms in SD2-FP and HR1-CH models. Such large atomic-scale calculation is truly unpreceded and can reveal many of the intricate details in bonding close to the location of the cleavage sites S1/S2 and S2’, discussed in the Introduction. As can be seen, mutations do slightly shift positions of the data points. However, details of specific bonding pairs close to the cleavage site is a daunting task. Most of the data points in **Figure 5(a)** with large BO values are from stronger covalent bonds. Those unaffected by mutations are strictly the internal covalent bonds within the AAs.

We would like to focus on the HBs with BL above 1.56 Å and the data are shown as star symbols. The strongest HB has BO of 0.12 e-at 1.56 Å. Mutation can either enhance or weaken the HB strengths. Even though HBs are relatively weak, they are ubiquitous and constitute a significant component of interatomic bonding in biomolecules (see **Figure 5(b)**). **Figure 5(b)** also shows data points of bonds at separations above 2.0 Å. There are four main groups of bonds: H-H, C-H, N…H and C-C indicated in the middle part of the **Figure 5(b)**. They all indicate that for accurate description on interatomic bonding and the effect of mutation, interactions for atomic pairs up to at least 4.5 Å is necessary.

**Figure 5.**
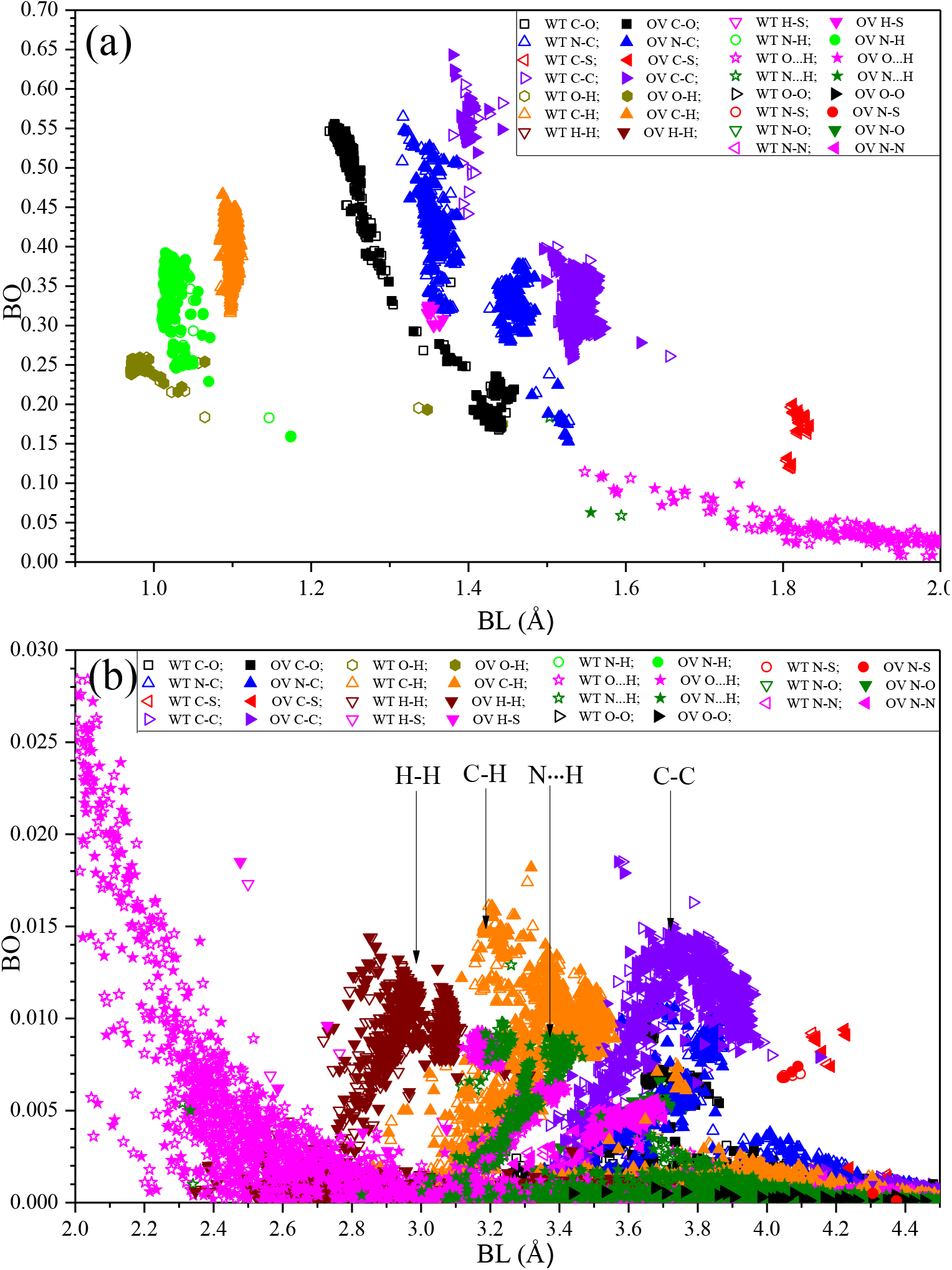
Overlap distributions of BO vs BL for the WT and OV SD2-FP and HR1-CH models with (a) 0.9 Å to 2.0 Å BL and (b) 2.0 Å to 4.5 Å. WT and OV are shown in open and closed symbols respectively.

In **Figure 5(a)** near 1.16 Å the covalent N-H bond has BO lowered by mutation. At 1.39 Å, the covalent O-H bond, mutation makes a negligible difference. At 1.52 Å, for covalent N-C bond mutation slightly lowered the BO. Near 1.63 Å, covalent C-C bond, mutation increased the BO. In **Figure 5(b)** at 2.49 Å, the covalent H-S bond increased BO due to mutation. For HBs in both **Figure 5(a)** and **(b)**, mutation can both increase and/or decrease the BO by a small amount. All this indicates that at the atomistic level the mutations can increase or decrease the bond strength, a situation correctively reflected in the column AABP (HB) of **Table 1**. Quantitatively, the total number of N…H and O…H HBs in the SD2-FP model for WT (OV) is 1285 (1285) and 2080 (2091) respectively, indicating an increase in O…H but no change in N…H. On the other hand, the number of HBs in the HR1-CH model for WT (OV) is 1039 (1063) for N…H and 1721 (1720) for O…H, showing a change in N…H but one less HB in O…H of OV. This analysis reveals that the intramolecular HBs alters slightly due to OV mutations.

### 2.4 Partial Charge

**Figure 6(a)** shows PC* distribution of the AABPU listed in **Table 1** for the 10 mutations. It is really striking to see that nearly all the PC* on Omicron are positive. The only mutation with negative PC* is H655Y. The other 9 mutations, or 90%, have positive PC* and all exhibiting a significantly increase in PC* after mutation. This a very important result obtained from our *ab initio* DFT calculation and has profound implications since the cell membranes in human body are mostly negatively charged [34]. Therefore, these mutated AAs will have enhanced non-specific electrostatic interactions with the cell membrane units. The shift toward positive PC* of OV mutations has a direct impact on the cleavage sites (indicated by vertical red dotted lines), especially the large increase in N679K and P691H, which are very close to the furin S1/S2 site and notable increase between D796Y and N856K that are surrounding the S’ site. This increase in the number of positive charges is crucial for the host protease in order to cleave the S-protein. However, several experimental studies have found that the Omicron S-protein cleaves less efficiently than other VOCs [12, 30–32], while another study has indicated the opposite [29]. This contradiction indicates that increased PC in Omicron may have other consequences, not yet elucidated. For example, N764K, N856K, N969K mutations have been observed to form interprotomer electrostatic contacts with the neighboring protomers, enhancing S-protein stability [35].

**Figure 6.**
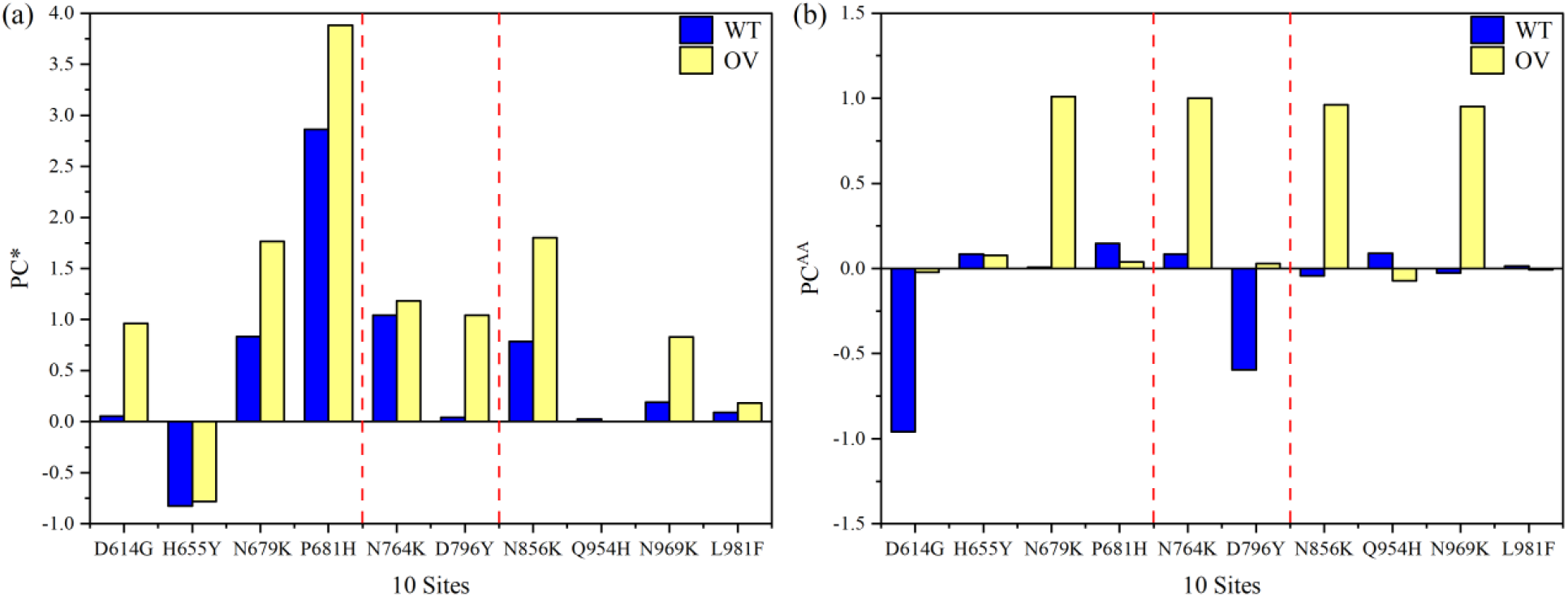
(a) Partial charge per AABPU for 10 mutations, WT (blue) and OV (yellow). The two dashed red lines shows the two cleavage sites. (b) PC^AA^ is the partial charge distribution of the AA at each unique site.

The trend toward positive charges of OV mutations is emphasized further in **Figure 6(b)**, detailing the PC^AA^ of the key AA at each site. To easy distinguish, we used two PC symbols: PC* for whole unit of AABPU and PC^AA^ for only one unique AA site. Interestingly, D614G, N679K, N764K, D796Y, N856K, and N969K OV mutations significantly shift their PC^AA^s to positive, whereas H655Y, Q954H, and L981F have minor or negligible changes. P681H exhibits a distinct behavior when comparing **Figures 6(a)** and **6(b)**, since the substitution of P to H changes the PC* of AABPU rather than the key AA itself. The reason could be connected to the fact that we performed our simulation at neutral pH, while the pH impact could play a significant role in the cleavage process, particularly regarding the charge of 681 site.

## 3. Discussion

### 3.1 AABPU as a specific structural unit

Protein-protein interactions (PPIs) are essential for many biological processes, so they have to be accurately identified not only at the molecular level but also at the AA and atomic levels. In fact, the AA-AA pairing governs these PPIs as well as protein folding and assembly. In this paper, we are strongly advocating the concept of AABPU, based on the AABP interaction proxy, as a special structural unit to describe and quantify atomistic level resolution of biomolecular structures and interactions. In particular, we apply it to S-protein domains in WT and OV cases. AABPU includes many key parameters that have not been emphasized by the scientific community such as interatomic interaction, PC, size, shape and so on. It goes beyond the traditional approach of using AA sequences only in analyzing and explaining most observations. AABPU may be useful for studying biological networks ranging from AA-AA pairing to 3D protein structure and protein-protein networks based on accurate *ab initio* approach. It provides new insights into the geometry alterations and interatomic interactions of different S-protein domains in WT and mutated type, significantly enhancing our understanding of the inter-and intra-molecular bonding between them as well as determining the effect of mutations arising from OV at specific S-protein domains and how they are impacting the cleavage sites. This concept could also be extended to investigate the mutational effects on other S-protein domains and S-protein protomers not covered in this study. However, there are also some major challenges that cannot be avoided in QM calculations rather than the AABPU concept. For example, the whole trimeric S-protein contains around 73000 atoms, which is currently impossible to calculate in a single *ab initio* simulation. In this regard, we have been developing a *divide and conquer strategy* [20, 21] to overcome this issue. But even with this development the simulations of whole domains of trimeric S-protein using QM methodologies are still computationally too expensive. Nonetheless, to our knowledge we believe that our simulation is the largest single *ab initio* calculation, performed on whole specific domains rather than AA fragments of a large biomolecular system such as the interface RBD-ACE2 complex, as described in the literature [36, 37]. Furthermore, this single AABPU quantifier has extensive parameters for gaining complementary microscopic knowledge of protein structure and protein-protein interactions, which could help us understand their features and functions better.

### 3.2 Role of cleavage sites in viral transmission of Omicron variant

The cleavage efficiency of S-protein at the furin-cleavage S1/S2 site increases the chances that the newly generated virus will be able to fuse rapidly with the host membrane. In this regard, mutations in the Alpha and Delta variants’ S1/S2 junction region have been shown to play an important role in improving the cleavage efficiency at this site [10,11]. Specifically, the Delta variant had a higher transmission rate than Alpha, owing to its P681R mutation vs P681H mutation of Alpha at the nearby location of the S1/S2 junction [28]. The reason for this could be due to the difference in the partial charge of H vs R, with the latter having a positive charge at all pH conditions whereas the former has a positive charge at mildly basic pH. Omicron, like Alpha, has a P681H mutation, so its cleavage efficiency may be less than that of the Delta variant, as recently reported [12]. Additionally, Omicron bears two more mutations close to this site, H655Y and N679K, which are likely to impact its S-protein cleavability by adding more positive charges to this site, but there are many debates about this prediction. Some studies have found that Omicron S-protein cleaves to be less effective than other variants [12,30–32], while others revealed the opposite view that the presence of the N679K with P681H mutations in Omicron significantly increases furin-mediated S1/S2 cleavability [29].

On the other hand, the cell-cell fusion requires another cleavage site, the S’ site. The protease activity can occur either by TMPRSS2 at the plasma membrane or cathepsins in the endosomes [7]. TMPRSS2 activity has been established to be critical for viral spread and pathogenesis in the infected host [7]. Indeed, it has been claimed that the TMPRSS2 is necessary for optimal virus entry in both the WT and Delta variants, but not in Omicron [12]. This suggests that Omicron prefers the endocytic cell entry route instead of the plasma membrane entry route [12,31,32]. This is consistent with the idea that reduced cleavage efficiency of Omicron S-protein at the S1/S2 site impairs TMPRSS2-mediated entry [12].

Together, the role of cleavage sites of Omicron is still unclear and requires further investigation. The main goal of this work is to speculate on the connection between Omicron mutations at proximal cleavage sites and the cleavability and fusogenicity, based on results from large-scale computation. It was clear from that the Omicron mutations inducing more positive partial charge distribution of S-protein. This change may play a function in controlling proteolytic cleavage, render S-protein more sensitive to low-pH induced conformational changes, promote an adaptation that utilizes the low-pH endosomal entry pathway, and/or boost virus entrance in the lower pH environment of the upper airway [12].

Unlike the Delta variant, the high transmissibility of OV may not be due to its efficient cleavability, but rather to the immune evasion and the strong binding affinity between RBD and ACE2 [6,12,30–32]. Therefore, more studies are needed to explore the impact of Omicron mutations on neutralizing antibodies and ACE2 binding, and how they are responding to other Omicron subvariants, like BA2 or BA3. We believe that the AABPU-based approach we developed can provide valuable information for identifying key epitopes on antibodies and in probing biological properties of Omicron subvariants or of any future VOC.

### 3.3 Towards Machine Leaning

Another important direction that has not been addressed above is how to use the calculated data on AABPU in Machine Leaning (ML) applications in a creative way. ML depends on the existence of large amount of data points to facilitate the reliable prediction. Additionally, ML should be feed with high-accuracy input data representing all S-protein variants that have occurred or are expected to occur in different domains to obtain accurate prediction. The data should be calculate based on the realistic 3D structure of the proteins, rather than the traditional approach of using only the primary sequence. We anticipate using our high-accuracy data on AABPU as input to ML algorithm. In fact, the data that we have generated so far or that will be calculated is unique and cannot be found elsewhere. Because of this, we will be limited to using only our data to verify the accuracy of ML-predicted data. However, it should be mentioned that large data set are necessary to predict the potential new variants as well as the effect of the specific mutation and their consequences. Currently, we are at the stage of collecting accurate unique data that will be provide insight knowledge about the interatomic interactions in 3D including implicitly all AA-AA network in S-protein. We expect our project will provide at least 200 data points including the 20 data points in **Table S1** for 10 mutations based on the data from **Table 1**. Importantly, the key step is to provide specific data points in the form that can be read by machine as shown in the last column of **Table S1**. Furthermore, **Table S2** shows our preliminary results for the 16 mutations in Omicron in RBD-SD2 domain [unpublished data]. The 56 data points from **Table S1** and **S2** enable us to start some preliminary tests such as using ¾ of the data to predict the other ¼. Our hopeful goal is using a smaller sample of highly accurate data based on our AABPU technique to predict the future mutations in S-protein. Of course, the success of this approach still needs to be tested and verified.

In conclusion, in this paper, we have:

1. developed a quantitative scheme and methodologies for rapid assessment of the effect of 10 mutations at the SD2-FP and HR1-CH domains of the Omicron variant.
2. connected bonding characteristic, size, and partial charge with to the existence of 2 cleavage sites S1/S2 and S2’ for the first time, showing that the Omicron mutations alter their physical and geometry features near to both sites.
3. conjectured that the possible reason for fast infection rate in Omicron variant is based on the bonding characteristic, size and partial charge as quantified by AABPU.

## 4. Material and Methods

### 4.1 Model construction

For large-scale *ab initio* quantum chemical calculation of a complex biomolecular system, the first step is to design the starting atomistic model. Based on the *divide and conquer strategy* [20,21], two such systems of SD2-FP and HR1-CH in unmutated and mutated forms are constructed as depicted in **Figure 1**. These systems contain 243 and 200 amino acids with 3654 and 3054 atoms for WT and 3681and 3671 atoms for mutated case. We followed the same approach, previously developed in the Delta variant simulation, to build these systems [21]. Briefly, the initial structure for SARS-CoV-2 S-protein shown in **Figure 1** was obtained from Woo *et al* from [PDB ID 6VSB] [38] which originated from Wrapp et al. study [17]. We chose Chain A in its up conformation and used to create the wild-type (WT) and mutated Omicron variant (OV) models. Sequence numbers for the SD2-FP model and HR1-CH model are 592-834 and 835-1034 in 6VSB 1_2_1 [39] respectively. This procedure is summarized as follows. First, we selected all residues of the SD2 and FP region to create the SD2-FP model (residue 592 to 834). Next, we removed the glycans and the associated hydrogen (H) atoms from the SD2-FP model and later added the H atoms using the Leap module in the AMBER package [40,41], which then yields the unmutated or wild type (WT) model. This WT model used as a template to generate the mutated OV model of SD2-FP with six substitutions D614G, H655Y, N679K, P681H, N764K, and D796Y using the Dunbrack backbone-dependent rotamer library [42] implemented by the UCSF Chimera package [43]. Similar steps were followed to prepare the WT and OV models for HR1-CH. The HR1-CH OV model contains four mutations: N856K, Q954H, N969K, and L981F. All four models, two WT of SD2-FP and HR1-CH and two relevant OV models, were first minimized with 500 steepest descent steps and 10 conjugate gradient steps using UCSF Chimera to avoid bad clashes of unrealistic close atomic pairs. These four initial atomistic models SD2-FP and HR1-CH are further optimized to sufficient accuracy using VASP (see next subsection). **Figure S3** shows the final atomic-scale structure in the models used in the calculation, like the ribbon structure depicted on the **Figure 1(d)**.

### 4.2 Vienna ab initio simulation package (VASP)

In this study, we used two well established density functional theory (DFT) based packages: The Vienna *ab initio* Simulations package (VASP) [44] and the orthogonalized linear combination of atomic orbital OLCAO) method [45], which will describe in next subsection. The combination of these two different DFT codes, VASP and OLCAO, has been successfully used in a variety of areas, including large biomolecules, organic or inorganic disordered systems, etc.[20– 23,46–51].

The initial structure derived from the protein data bank (PDB) with appropriate modification and addition of hydrogen (H) atoms are fully relaxed by using VASP [44] known for its efficiency in structural optimization. The projector augmented wave (PAW) method with Perdew-Burke-Ernzerhof (PBE) exchange correlation functional [52] within the generalized gradient approximation (GGA) is one of the option we chose that balance the accuracy needed and the computational resources available. Detailed tests in the past suggest that the use of the following input parameters for biomolecular systems: (1) energy cut-off at 500 eV; (2) electronic convergence of 10^−4^ eV; (3) force convergence for ionic steps at −10^−2^ eV/Å; (4) single k-point sampling at the center of the supercell. The final relaxed structure of the models has achieved with total energy difference less than −0.330 eV or −0.000108 eV per atom and will be used as the input for OLCAO calculations.

### 4.3 Orthogonalized linear combination of atomic orbitals (OLCAO) method

A different DFT method, the OLCAO method developed in-house [50], used to calculate the electronic structure and interatomic interactions of biomolecule systems. The OLCAO method uses minimal atomic basis expansion and orthogonalization to the core orbitals technique in order to deal with the huge matrix with a single diagonalization to obtain all the energy states and wave functions of the Kohn-Sham equation [45]. The key feature of the OLCAO method is the provision of two fundamental structural parameters: the effective charge (*Q*^∗^) on each atom and the bond order (BO) values *ρ*_*αβ*_ between any pair atoms αand βin the large supercell of the biomolecule. They are obtained from the *ab initio* wave functions with atomic basis expansion calculated quantum mechanically for the large biomolecule. The two OLCAO structural parameters are defined as

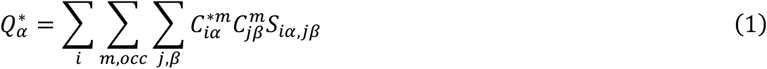

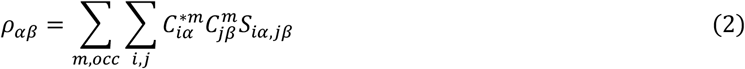

In the above equations, *S*_*iα,jβ*_ are the overlap integrals between the *i*^*th*^ orbital in the *α*^*th*^ atom and the *j*^*th*^ orbital in the *β*^*th*^ atom. 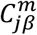 are the eigenvector coefficients of the *m*^*th*^occupied molecular orbital. The deviation of 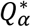 from the neutral atomic charge 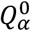 on the same atom *α* is usually referred to as the partial charge (PC) or 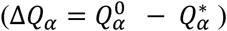 The BO quantifies *the strength of the bond between two atoms* in the unit of electrons (e^-^) and usually scales with the bond length (BL) or the distance of separation between atoms α and β,depending also on the configuration of the vicinal atoms. It should be pointed out that the calculated PC and BO in Eq. (1) and (2) is basis-dependent since they are based on the Mulliken scheme [53,54] using localized atomic orbitals. We use the minimal basis in all our calculations for biomolecular systems. Another very important point is that we use a one-point calculation to obtain all BO values for all atomic pairs to characterize the internal cohesion of the system under study, rather than the traditional total energy or enthalpy calculation (two-point or even many-point calculation) which is used to describe the strength of binding between biomolecular systems. The sum of all BO values within a structural component - such as a protein - gives the total bond order (TBO), which accurately describes the interatomic interactions that are critical to analyzing the AA-AA network for large complex biomolecules.

In complex biomolecules one can extend the concept of the bond order (BO), defined for a *pair of atoms*, and apply it to the interaction between a *pair of amino acids* (AAs). We refer to this generalized quantifier of molecular interactions the *amino acid-amino acid bond pair* (AABP) [23], defined as

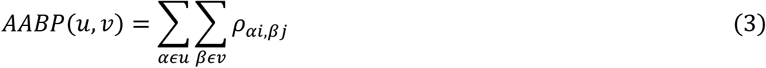

In Eq. (3), the summations are over all atoms *α* in *AA u* and all atoms *β* in *AA v*. AABP considers all possible bonding between two AAs including both covalent and hydrogen bonding (HB). AABP value is a single parameter proxy that quantifies the interaction between two AAs. The stronger the interaction, the higher will be the AABP and vice versa. AABP can be further resolved in nearest neighbor (NN-AABP) and non-local (NL-AABP) parts, which discussed in detail in **Section 2.1**. It should be emphasized that the AABP does not involve the “BL” used for the description of interacting atoms since the distance of separation between two AAs is difficult to quantify precisely. AABP values are calculated from quantum mechanical wave functions of the entire biomolecular system and thus represent a collective structural parameter, including the effects of all atomic pairs involved. AABP is a new fundamental measure of molecular interactions in biomolecules, that contains the nearest-neighbor or local interactions of AAs that are vicinal along the sequence and in the 3D folding space, as well as the off-diagonal or non-local interaction between AAs that are not vicinal in the sequence space but are interacting in 3D folding space.

## Data Availability

All calculated data is for the Figures and Tables is available upon request.

## Authors’ Contributions

WC, PA and BJ conceived the project. WC and PA performed the calculations. PA and BJ made most of the figures. BJ searched most of the references, WC, BJ, PA drafted the paper with inputs from RP. All authors participated in the discussion and interpretation of the results. All authors edited and proofread the final manuscript.

## Acknowledgements

This research used the resources of the National Energy Research Scientific Computing Center supported by DOE under Contract No. DE-AC03-76SF00098 and the Research Computing Support Services (RCSS) of the University of Missouri System. We thank Dr. Richard Gerber, Senior Science Advisor and HPC Department Head for special allocations. This project is funded partly by the National Science Foundation of USA: RAPID DMR/CMMT-2028803. RP acknowledges funding from the Key project #12034019 of the National Natural Science Foundation of China.

## Supplementary Materials

**Figure S1**. Ball and stick figures of SD2-FP and HR1-CH showing the two cleavage sites marked in magenta and Omicron variants mutations in the region of SD2-FP and HR1-CH are marked in green and orange respectively. Red: O, blue: N, grey: C, white: H, yellow. **Figure S2**. TDOS of the **(a)** SD2-FP model, **(b)** HR1-CH model for both WT and OV. **Figure S3**. PDOS of the WT and OV amino acids for six sites of **(a)** SD2-FP model, and four sites of **(b)** HR1-CH model. **Table S1**: Data notation in machine readable format for ML based on **Table 1**. The last digit is ‘0’ for WT and ‘1’ for the mutated type. **Table S2**: Data notation in machine readable format for 16 mutations in RBD-S2 domain (unpublished). *(Supplementary Materials)*

