## Supplementary Information for "Atomic-scale Quantum Chemical Calculation of Omicron Mutations Near Cleavage Sites of the Spike Protein"

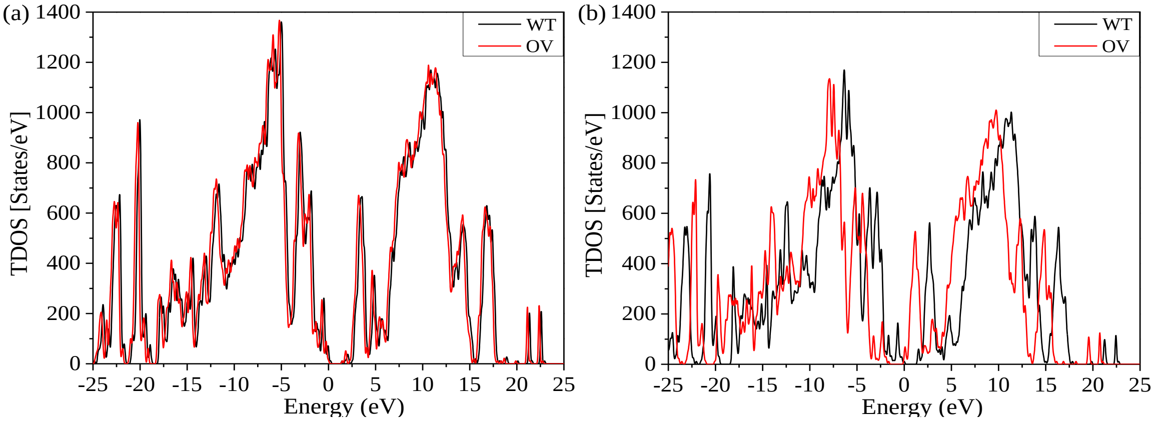


**Figure S1.** TDOS of the **(a)** SD2-FP model, **(b)** HR1-CH model for both WT and OV.

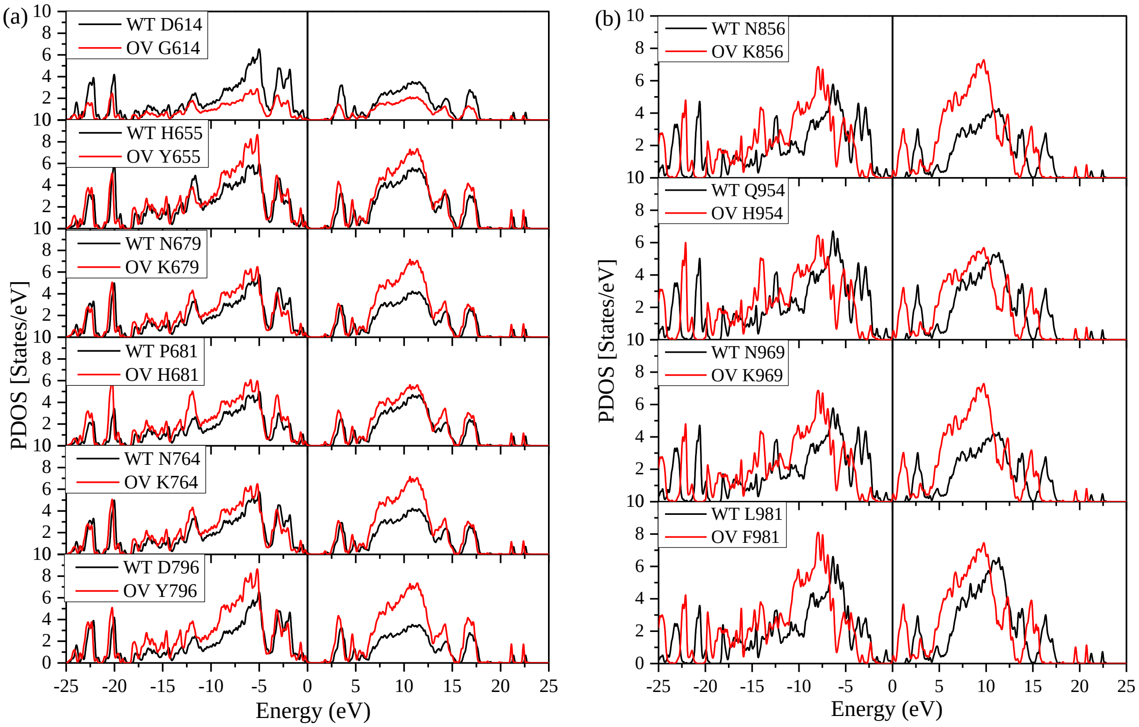


**Figure S2.** PDOS of the WT and OV amino acids for six sites of **(a)** SD2-FP model, and four sites of **(b)** HR1-CH model.

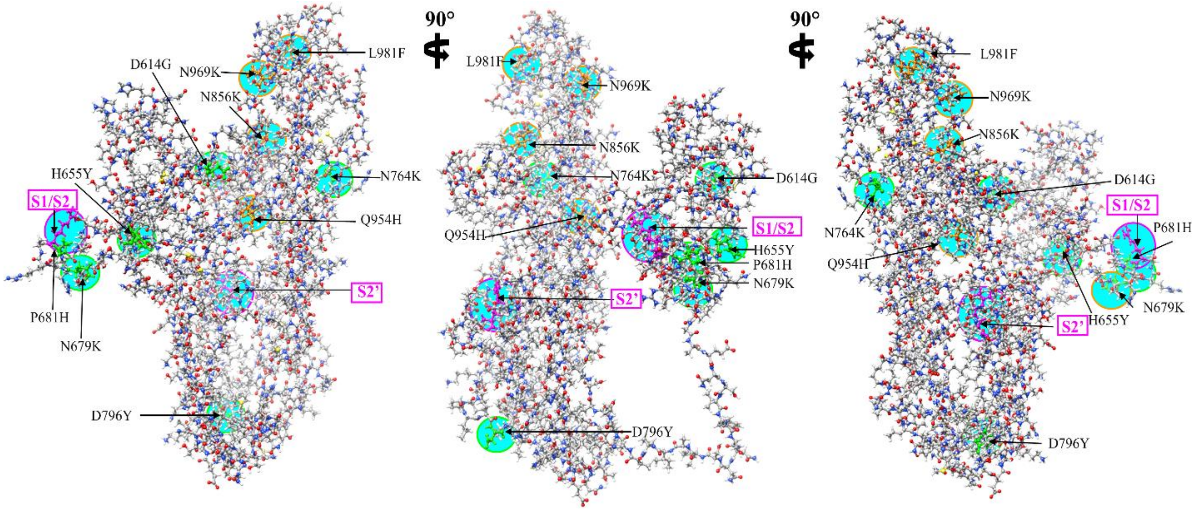


**Figure S3**. Ball and stick figures of SD2-FP and HR1-CH showing the two cleavage sites marked in magenta and Omicron variants mutations in the region of SD2-FP and HR1-CH are marked in green and orange respectively. Red: O, blue: N, grey: C, white: H, yellow: S.

**Table S1.** Data notation in machine readable format for ML based on **Table 1**. The last digit is '0' for WT and '1' for the mutated type.

| Models | Total AABP | NN-AABP | NL-AABP | AABP (HB) | Data notation |
|---|---|---|---|---|---|
| WT D614 | 0.917 | 0.912 | 0.005 | 0.040 | D614-0.917-0.912-0.005-0.040-0 |
| OV G614 | 0.908 | 0.907 | 0.002 | 0.042 | G614-0.908-0.907-0.002-0.042-1 |
| WT H655 | 0.976 | 0.968 | 0.007 | 0.032 | H655-0.976-0.968-0.007-0.032-0 |
| OV Y655 | 0.971 | 0.965 | 0.006 | 0.032 | Y655-0.971-0.965-0.006-0.032-1 |
| WT N679 | 1.022 | 0.956 | 0.066 | 0.081 | N679-1.022-0.956-0.066-0.081-0 |
| OV K679 | 1.011 | 0.963 | 0.048 | 0.072 | K679-1.011-0.963-0.048-0.072-1 |
| WT P681 | 1.117 | 1.064 | 0.054 | 0.064 | P681-1.117-1.064-0.054-0.064-0 |
| OV H681 | 1.032 | 0.984 | 0.048 | 0.068 | H681-1.032-0.984-0.048-0.068-1 |
| WT N764 | 1.130 | 1.008 | 0.121 | 0.137 | N764-1.130-1.008-0.121-0.137-0 |
| OV K764 | 1.118 | 1.019 | 0.100 | 0.118 | K764-1.118-1.019-0.100-0.137-1 |
| WT D796 | 1.175 | 1.124 | 0.052 | 0.066 | D796-1.175-1.124-0.052-0.066-0 |
| OV Y796 | 1.051 | 1.000 | 0.051 | 0.070 | Y796-1.051-1.000-0.051-0.070-1 |
| WT N856 | 0.935 | 0.894 | 0.041 | 0.069 | N856-0.935-0.894-0.041-0.069-0 |
| OV K856 | 0.937 | 0.902 | 0.036 | 0.065 | K856-0.937-0.902-0.036-0.065-1 |
| WT Q954 | 1.148 | 1.008 | 0.140 | 0.152 | Q954-1.148-1.008-0.140-0.152-0 |
| OV H954 | 1.146 | 1.008 | 0.139 | 0.155 | H954-1.146-1.008-0.139-0.155-1 |
| WT N969 | 0.938 | 0.907 | 0.031 | 0.052 | N969-0.938-0.907-0.031-0.052-0 |
| OV K969 | 0.946 | 0.913 | 0.033 | 0.053 | K969-0.946-0.913-0.033-0.053-1 |
| WT L981 | 0.898 | 0.893 | 0.005 | 0.036 | L981-0.898-0.893-0.005-0.036-0 |
| OV F981 | 0.917 | 0.888 | 0.029 | 0.059 | F981-0.917-0.888-0.029-0.059-1 |

**Table S2.** Machine-readable data notation in ML for the RBD-SD2 OV model (unpublished data). The last digit is '0' for WT and '1' for the mutated type.

| Models | Total AABP | NN-AABP | NL-AABP | AABP (HB) | Data notation |
|---|---|---|---|---|---|
| WT G339 | 1.016 | 0.993 | 0.023 | 0.052 | G339-1.016-0.993-0.023-0.052-0 |
| OV D339 | 1.196 | 1.154 | 0.042 | 0.063 | D339-1.196-1.154-0.042-0.063-1 |
| WT S371 | 0.918 | 0.888 | 0.030 | 0.051 | S371-0.918-0.888-0.030-0.051-0 |
| OV L371 | 0.945 | 0.928 | 0.017 | 0.040 | L371-0.945-0.928-0.017-0.040-1 |
| WT S373 | 0.941 | 0.920 | 0.021 | 0.052 | S373-0.941-0.920-0.021-0.052-0 |
| OV P373 | 0.999 | 0.992 | 0.008 | 0.031 | P373-0.999-0.992-0.008-0.031-1 |
| WT S375 | 0.944 | 0.916 | 0.028 | 0.058 | S375-0.944-0.916-0.028-0.058-0 |
| OV F375 | 0.926 | 0.917 | 0.009 | 0.037 | F375-0.926-0.917-0.009-0.037-1 |
| WT K417 | 1.216 | 1.013 | 0.203 | 0.203 | K417-1.216-1.013-0.203-0.203-0 |
| OV N417 | 1.066 | 1.017 | 0.048 | 0.069 | N417-1.066-1.017-0.048-0.069-1 |
| WT N440 | 0.985 | 0.981 | 0.005 | 0.037 | N440-1.066-0.981-0.005-0.037-0 |
| OV K440 | 0.983 | 0.978 | 0.005 | 0.037 | K440-0.983-0.978-0.005-0.037-1 |
| WT G446 | 0.912 | 0.910 | 0.002 | 0.038 | G446-0.912-0.910-0.002-0.038-0 |
| OV S446 | 1.038 | 0.979 | 0.059 | 0.091 | S446-1.038-0.979-0.059-0.091-1 |
| WT S477 | 0.964 | 0.958 | 0.006 | 0.039 | S477-0.964-0.958-0.006-0.039-0 |
| OV N477 | 1.157 | 1.156 | 0.001 | 0.151 | N477-1.157-1.156-0.001-0.151-1 |
| WT T478 | 1.044 | 1.043 | 0.001 | 0.022 | T478-1.044-1.043-0.001-0.022-0 |
| OV K478 | 1.214 | 1.212 | 0.002 | 0.139 | K478-1.214-1.212-0.002-0.139-1 |
| WT E484 | 1.040 | 0.927 | 0.114 | 0.124 | E484-1.040-0.927-0.114-0.124-0 |
| OV A484 | 0.934 | 0.932 | 0.002 | 0.030 | A484-0.934-0.932-0.002-0.030-1 |
| WT Q493 | 1.060 | 0.973 | 0.087 | 0.106 | Q493-1.060-0.973-0.087-0.106-0 |
| OV R493 | 1.165 | 1.165 | 0.194 | 0.200 | R493-1.165-1.165-0.194-0.200-1 |
| WT G496 | 0.975 | 0.944 | 0.031 | 0.062 | G496-0.975-0.944-0.031-0.062-0 |
| OV S496 | 0.994 | 0.938 | 0.055 | 0.076 | S496-0.994-0.938-0.055-0.076-1 |
| WT Q498 | 1.120 | 1.073 | 0.047 | 0.054 | Q498-1.120-1.073-0.047-0.054-0 |
| OV R498 | 1.179 | 1.056 | 0.123 | 0.126 | R498-1.179-1.056-0.123-0.126-1 |
| WT N501 | 1.120 | 1.073 | 0.047 | 0.054 | N501-1.120-1.073-0.047-0.054-0 |
| OV Y501 | 1.034 | 0.942 | 0.092 | 0.104 | Y501-1.034-0.942-0.092-0.104-1 |
| WT Y505 | 1.058 | 0.974 | 0.084 | 0.104 | Y505-1.058-0.974-0.084-0.104-0 |
| OV H505 | 0.998 | 0.953 | 0.045 | 0.069 | H505-0.998-0.953-0.045-0.069-1 |
| WT T547 | 1.033 | 0.977 | 0.056 | 0.079 | T547-1.033-0.977-0.056-0.079-0 |
| OV K547 | 0.994 | 0.977 | 0.016 | 0.042 | K547-0.994-0.977-0.016-0.042-1 |
